# Diversity of microglial transcriptional responses during opioid exposure and neuropathic pain

**DOI:** 10.1101/2021.12.29.473853

**Authors:** Elizabeth I Sypek, Hannah Y Collins, William M McCallum, Alexandra T Bourdillon, Ben A Barres, Christopher J Bohlen, Grégory Scherrer

## Abstract

Microglia take on an altered morphology during chronic opioid treatment. This morphological change is broadly used to identify the activated microglial state associated with opioid side effects, including tolerance and opioid-induced hyperalgesia (OIH). Following chronic opioid treatment and peripheral nerve injury (PNI) microglia in the spinal cord display similar morphological responses. Consistent with this observation, functional studies have suggested that microglia activated by PNI or opioids engage common molecular mechanisms to induce hypersensitivity. Here we conducted deep RNA sequencing of acutely isolated spinal cord microglia from male mice to comprehensively interrogate transcriptional states and mechanistic commonality between multiple OIH and PNI models. Following PNI, we identify a common early proliferative transcriptional event across models that precedes the upregulation of histological markers of activation, followed by a delayed and injury-specific transcriptional response. Strikingly, we found no such transcriptional responses associated with opioid-induced microglial activation, consistent with histological data indicating that microglia number remain stable during morphine treatment. Collectively, these results reveal the diversity of pain-associated microglial transcriptomes and point towards the targeting of distinct insult-specific microglial responses to treat OIH, PNI, or other CNS pathologies.

## Introduction

Microglia, the resident macrophages of the central nervous system (CNS), regulate a host of neural processes such as synaptic pruning and programmed cell death ^1,2^. In settings of injury or disease, microglia enter an activated state characterized by changes in number, morphology, and production of signaling molecules ^3,4^. Following peripheral nerve injury, microglia morphology and distribution evolve in the spinal cord dorsal horn, exhibiting shortened processes and increased density where injured sensory neuron terminate ^5–7^. These observations have led to numerous functional studies that established the contribution of activated microglia to the development of chronic neuropathic pain ^8,9^.

We and others have observed a similar activated morphology following chronic opioid treatment, corresponding with the onset of tolerance and opioid-induced hyperalgesia (OIH) ^10–12^. Given these shared morphological alterations, common molecular mechanisms are thought to underlie the deleterious pronociceptive functions of microglia for both opioid- and injury-induced pain states. The ATP receptor P2X_4_ is upregulated in spinal microglia after both peripheral nerve injury and chronic morphine treatment, and P2X_4_ signaling is required for the development of tactile hypersensitivity after both spinal nerve injury and morphine tolerance ^13,14^. Both neuropathic pain and OIH have been suggested to rely on P2X_4_-mediated release of BDNF from microglia. BDNF could then act on TrkB expressed by neurons to downregulate the chloride transporter KCC2 and disturb chloride homeostasis, resulting in pronociceptive neuronal hyperexcitability in the spinal cord dorsal horn ^11,13,15^. In addition, P2X_7_, another microglial purinergic receptor, is upregulated and implicated in both morphine tolerance ^13,16–18^ and neuropathic pain ^19^. While we found no evidence of expression of the mu opioid receptor (MOR) or its mRNA (*Oprm1*) in microglia ^10^, others have suggested that opioids can act directly on microglia toll-like receptor 4 (TLR4) ^20,21^, and this receptor may also contribute to neuropathic pain ^22^. Further, we reported that morphine-induced microglial reactivity is intact in global MOR knockout mice, but that these animals do not display OIH ^10^. These results suggested a mechanistic dissociation between microglia activation and the function of these cells in OIH, despite previous evidence that inhibiting microglial reactivity can prevent morphine side effects ^23^.

While many potential molecular mechanisms of microglial activation in pronociceptive states have been identified, a complete understanding of microglial responses across pronociceptive conditions is required to determine if microglial activation-induced pain facilitation is a singular phenomenon. A clearer understanding of glial response to pain states could identify selective interventions that precisely alter microglia function and reduce hyperalgesia. The ability to isolate individual cell types and sequence their RNA to resolve their transcriptome has drastically expanded our understanding of cell-specific roles and intercellular interactions. With these methods, we can compare microglia transcriptomes to those of other CNS cell types ^24^, immune cells ^25–27^, and even across disease states ^28–31^ and brain regions ^32^. However, there has been no systematic comparison of microglial responses across pain states and chronic opioid treatment to directly ask whether injuries and opioids engage similar microglial pronociceptive mechanisms.

To address this issue, we set out to use transcriptomics to resolve microglial response patterns in multiple settings of heightened pain sensitivity. This analysis commands a reappraisal of the microglial mechanisms that underlie injury- and opioid-induced hypersensitivity and a redefinition of microglia activation based on transcriptional rather than histological profiles.

## Results

### Morphine Exposure does not induce strong transcriptional responses in microglia

We and others have previously shown histological evidence of microglia activation following chronic morphine treatment that coincides with the establishment of tolerance and OIH ^10^. Moreover, this activation is intact in mu opioid receptor (MOR) knockout mice and thus independent of MOR signaling ^10^. We therefore used transcriptomics to clarify the MOR-independent molecular mechanisms associated with opioid-induced microglial reactivity. We first treated mice with 10mg/kg morphine for 10 days, a dosing paradigm known to produce tolerance, OIH, and microglial activation (Figure 1A, B). We then isolated the lumbar spinal cord, as this region was particularly important for further comparisons to sciatic nerve injury conditions. We combined mechanical dissociation of tissue and magnetic activated cell sorting (MACS) for myelin depletion and CD11b selection to isolate a highly enriched microglial population for transcriptional analyses (Figure 1C). As we relied on CD11b surface expression to enrich for microglia, we also captured a small number of other myeloid cells such as neutrophils and perivascular macrophages. However, we saw strong enrichment of microglial genes compared with other non-myeloid CNS cell types including endothelial cells, oligodendrocytes, neurons and astrocytes (Figure 1D). Note we did not find evidence of infiltration or expansion of monocytes and neutrophils in any chronic pain model, consistent with previous studies ^33–35^.

**Figure 1:**
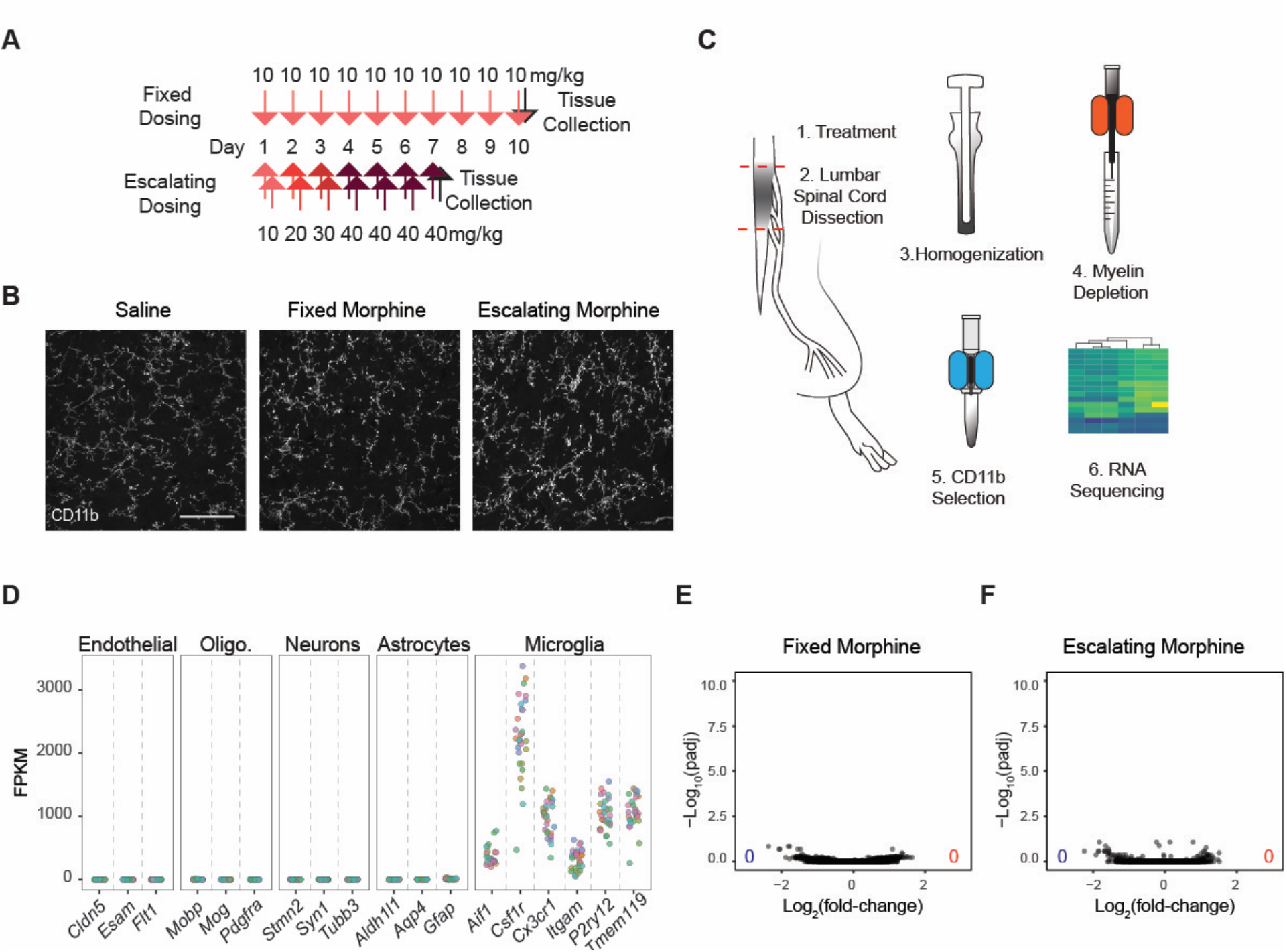
Minimal transcriptomic changes occur with chronic morphine-induced microglial activation. (A) Schematic illustrating the two opioid dosing paradigms. Fixed dosing was once daily 10mg/kg (subcutaneous, s.c.) for 10 days, with tissue collection 1 hour after the final dose on day 10. Escalating dosing is twice daily dosing of 10, 20, and 30 mg/kg per day (s.c.) for the first three days, then 3 days of twice daily 40mg/kg, and a final day of a single dose of 40mg/kg 1 hour before perfusion. (B) Example images of CD11b immunoreactivity of spinal cord dorsal horn after saline treatment, fixed morphine, or escalating morphine treatment. Scale bar = 50 μm. (C) Schematic of location tissue collection and microglia isolation for RNA sequencing (RNA-seq). (D) Normalized reads for genes specifically expressed in possibly contaminating cell types and markers of microglia, from all conditions and replicates tested. FPKM = Fragments Per Kilobase of transcript per Million mapped reads. (E and F) Volcano plots summarizing gene expression changes in microglia after fixed (E) or escalating (F) morphine treatment. Blue and red symbols indicate significantly down or upregulated genes from Deseq2, respectively (padj<0.01). Triangles indicate values outside the axis scale. The numbers in the lower corners indicate the total numbers of down (blue) and upregulated (red) genes (padj<0.01).

Strikingly, despite obvious morphological changes suggesting an activated state (Figure 1B), chronic morphine treatment induced no significant changes in microglial gene expression (Figure 1E). This is in stark contrast to hundreds of genes showing differential expression in other chronic pain models when tested using the same methods (see below). We interrogated this lack of transcriptional response further with a more aggressive dosing schedule, an escalating dosing paradigm that produces even greater histological evidence of activation, and again detected no transcriptional variation compared to control (Figure 1A,B,F).

We next used an *in vitro* culture system to further examine these unexpected observations. Microglial cultures have been used extensively to interrogate microglial response to opioids ^16,33,34^, and some studies have suggested that morphine stimulation of TLR4 may contribute to microglial activation^35^. To investigate the direct effects of morphine on microglia, we utilized a highly pure serum-free culture system ^36^ to compare morphine treatment to TLR2 and TLR4 agonists (LPS, PAM3CSK4, and zymosan) known to induce robust transcriptional changes. As expected, all three TLR agonists produced extensive gene expression changes, but again we did not observe any transcriptional changes with morphine treatment (Figure 2A-D). To confirm this finding, we conducted additional treatments of cultures with LPS or morphine (1uM up to 10uM) and performed qPCR on genes associated with TLR activation. Again, significant gene upregulation was detected with LPS but not with any morphine concentrations (Figure 2E). To ensure that the high variability between TLR agonism and control did not mask potential differential expression in morphine treatment, we again ran Deseq2 to compare morphine treatment to PBS alone and still did not identify any differences. Note that consistent with our previous reported ^10^, we did not detect *Oprm1* transcripts in serum free cultured microglia, either before or after morphine or TLR agonist exposure. Potentially, morphine could activate microglia following binding to the other opioid receptor type subtypes delta (DOR) or kappa (MOR). To test this possibility, we examined our transcriptional datasets and found no transcript for *Oprd1* or *Oprk1*, suggesting that canonical opioid signaling does not occur in microglia.

**Figure 2:**
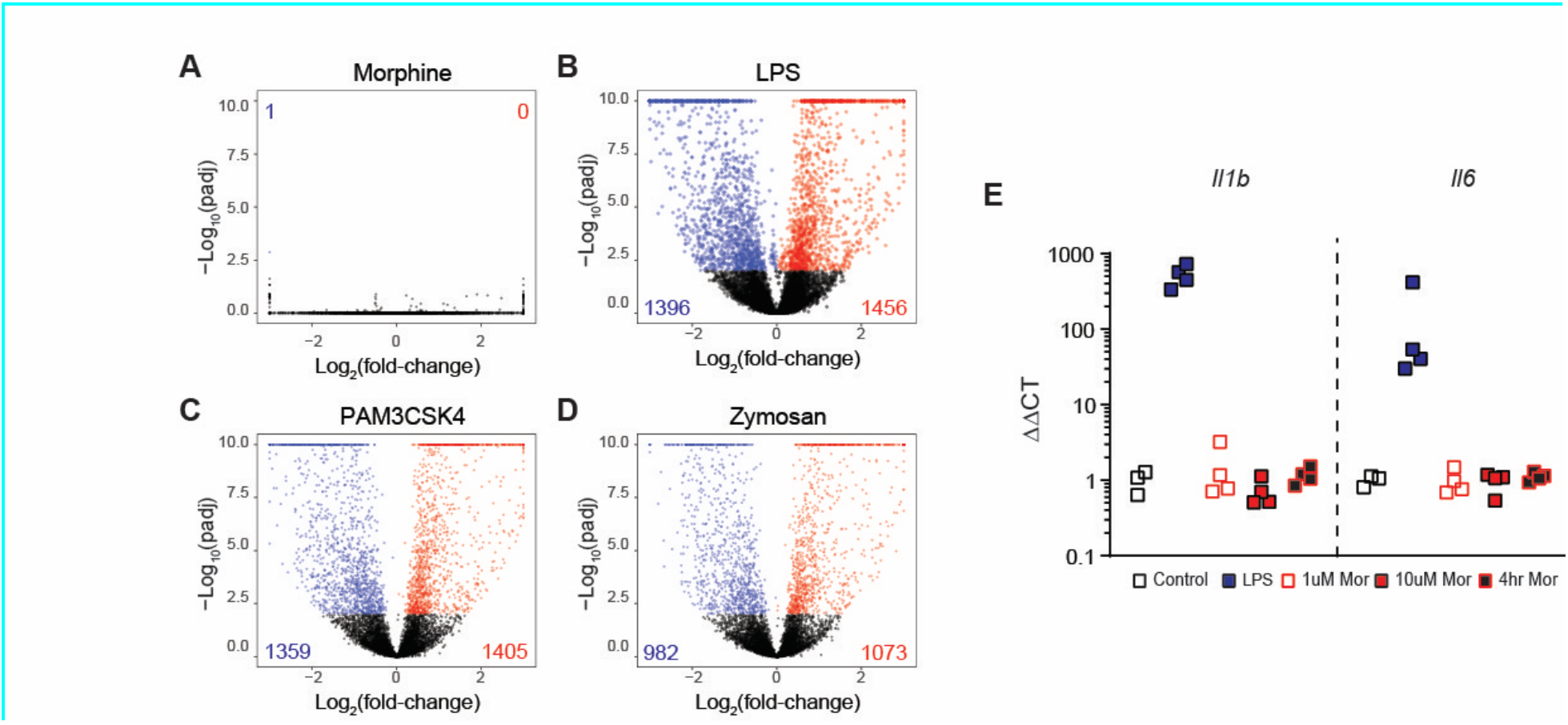
Morphine treatment differs from TLR agonism in cultured microglia. (A-D) Volcano plots summarizing gene expression changes in cultured rat cortical microglia after 24hr of treatment with 1uM morphine (A), LPS (1 ng/mL) (B), PAM3CSK4 (1 ng/mL) (C), or Zymosan (100 ug/mL) (D). Blue and red symbols indicate significantly down or upregulated genes from Deseq2, respectively (padj<0.01). Triangles indicate values outside the axis scale. The numbers in the lower corners indicate the total numbers of down (blue) and upregulated (red) genes at padj<0.01. (E) qPCR for *Il1b* and *Il6* 24 hours after LPS or morphine (1 or 10uM) exposure and 4 hours post 10uM morphine.

We conclude from both our *in vivo* and *in vitro* experiments that morphine exposure has minimal impact on microglial gene expression profiles, even at doses that alter microglial morphology *in vivo* and produce behavioral hypersensitivity.

### Transcriptional response to PNI differs across injury types

Given the reported mechanistic similarities of microglia activation and pronociceptive action following PNI, we next directly compare chronic opioid treatment and PNI models on a transcriptional level. We used the sequencing approach described above to uncover the diversity of transcriptional changes that occur in microglia in response to multiple peripheral injury models broadly used in the pain field. First, we used the spared nerve injury (SNI) model ^37^, which consists of transecting two of the three branches of the sciatic nerve (Figure 3A). As expected, SNI induced a strong increase in CD11b immunoreactivity 7 days post injury (dpi) when comparing ipsilateral to contralateral spinal cord, indicating microglial activation (Figure 3B). Remarkably, however, differential expression analysis with DESeq2 revealed only minimal gene expression changes in isolated microglia 7 dpi (adjusted p value (padj) <0.01), with only 7 downregulated and 1 upregulated gene, the complement pathway component *C4b* (Figure 3C).

**Figure 3:**
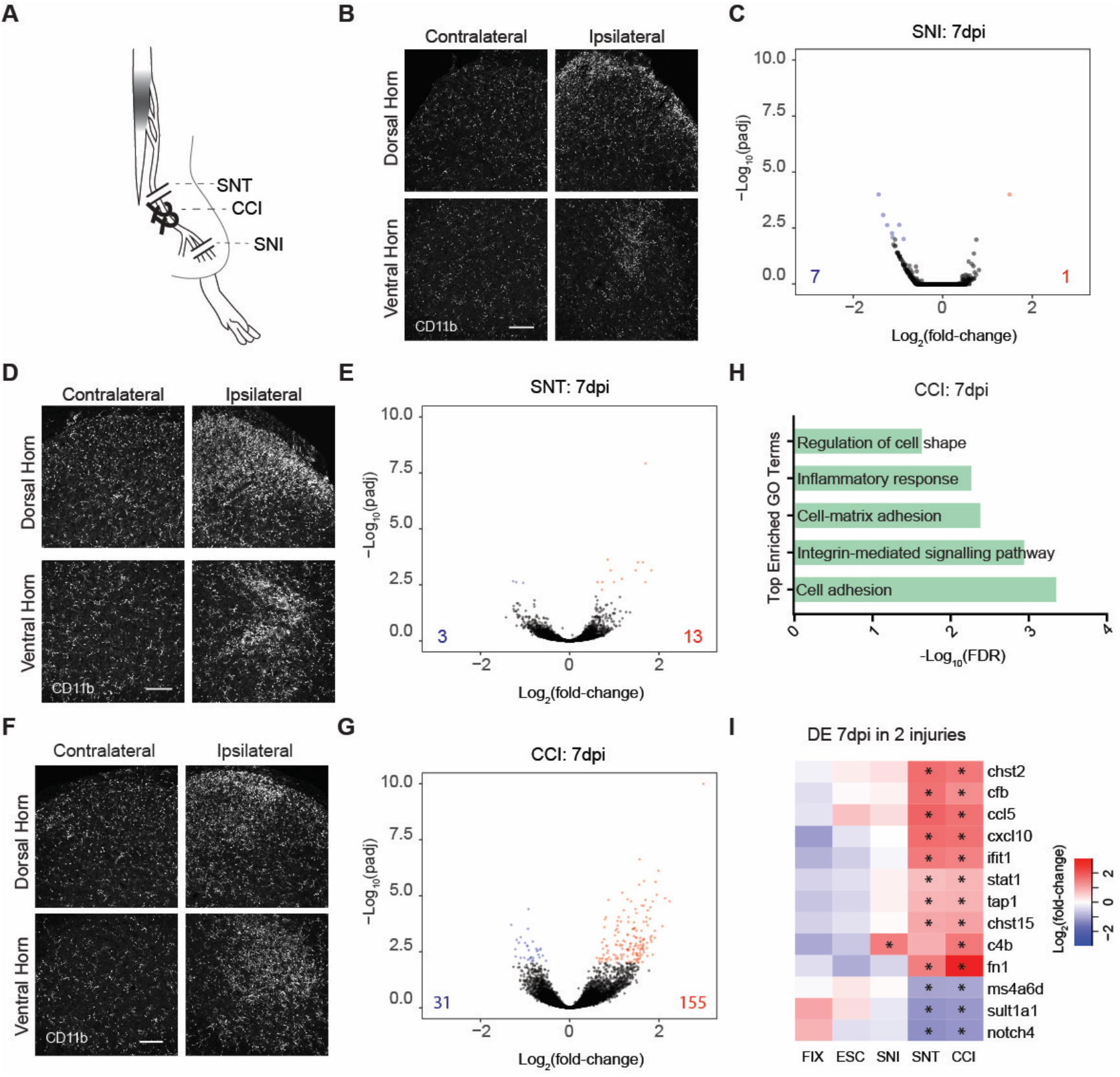
Diverse transcriptional response of spinal microglia to chronic neuropathic pain conditions. (A) Schematic of peripheral nerve injuries: complete sciatic nerve transection (SNT), chronic constriction injury (CCI) and spared nerve injury (SNI). (B) Example images of Cd11b immunostaining in lumbar spinal cords 7 days after unilateral SNI. dpi=days post injury. Scale bar = 100 μm. (C) Volcano plots summarizing gene expression changes in microglia 7 days after bilateral SNI. Blue and red symbols indicate significantly down or upregulated genes from Deseq2, respectively (padj<0.01). Triangles indicate values outside the axis scale. The numbers in the lower corners indicate the total numbers of down (blue) and upregulated (red) genes at padj<0.01. (D) Example images of CD11b immunostaining in lumbar spinal cords 7 days after unilateral SNT. dpi=days post injury. Scale bar = 100 μm. (E) Volcano plots summarizing gene expression changes in microglia 7 days after bilateral SNT. Blue and red symbols indicate significantly down or upregulated genes from Deseq2, respectively (padj<0.01). Triangles indicate values outside the axis scale. The numbers in the lower corners indicate the total numbers of down (blue) and upregulated (red) genes at padj<0.01. (F) Example images of CD11b immunoreactivity in the spinal cord 2 and 7 days after unilateral CCI. Scale bar = 100 μm. (G) Volcano plots summarizing gene expression changes in microglia 7 (D) days after bilateral CCI. Blue and red symbols indicate significantly down or upregulated genes from Deseq2, respectively (padj<0.01). Triangles indicate values outside the axis scale. The numbers in the lower corners indicate the total numbers of down (blue) and upregulated (red) genes at padj<0.01. (H) Top enriched GO Terms associated with CCI 7 days post injury. (I) Heatmap of genes differentially regulated (padj<0.01 in Deseq2) in both transection and CCI or SNI and CCI at 7dpi plotted as Log_2_(fold-change). Stars indicate significance in that condition.

As SNI leaves a large proportion of somatosensory neurons intact ^6^, we hypothesized that the low detection of SNI-induced differential RNA expression could result from a dilution effect associated with the presence of resting state microglia. To test this possibility, we used a more severe injury, a complete bilateral sciatic nerve transection (SNT) (Figure 3A). Immunohistological analysis of mice that underwent unilateral SNT again confirmed previous reports of marked microglial activation (Figure 3D). As with SNI, and despite this substantial change in CD11b staining pattern, quantification of differential gene expression supported our previous results that nerve injury-induced microglial activation is accompanied by minimal gene expression changes 7 days after injury; we found that only 3 genes were significantly downregulated, while 13 were upregulated (padj<0.01, Figure 3E).

### Selective and persistent microglial immune response after CCI

Having observed minimal transcriptional changes 7 days after full or partial transection of the sciatic nerve, we next utilized a third allodynia-inducing peripheral nerve injury model: chronic constriction injury (CCI) ^38^. In this model, two ligatures are loosely tied around the full thickness of the sciatic nerve (Figure 3A). CCI caused a similar pattern of CD11b immunoreactivity as in SNT and SNI, with an increase in CD11b expression at 7dpi (Figure 3F). However, specifically in this injury condition, a much larger number of differentially expressed genes were detected, with 31 downregulated genes and 155 upregulated genes (Figure 3G), uncovering a profound divergence of microglial responses between different peripheral nerve injuries. To characterize this unique response to CCI, we utilized Gene Ontogeny (GO) term analysis and found that the microglial response 7 days after chronic constriction injury was predominately related to immune and inflammatory processes (Figure 3H), including members of the complement pathway (*C4b* and *Cfb*) and cytokines (*Ccl5* and *Cxcl10*). Among the 13 genes upregulated at 7 days post full transection, 9 were also upregulated with CCI at this time point, most of which being implicated in immune responses, including *Cfb, Ifit1*, and *Ccl5* (Figure 5I) while overlap between gene expression changes in CCI and SNI was much lower.

**Figure 4:**
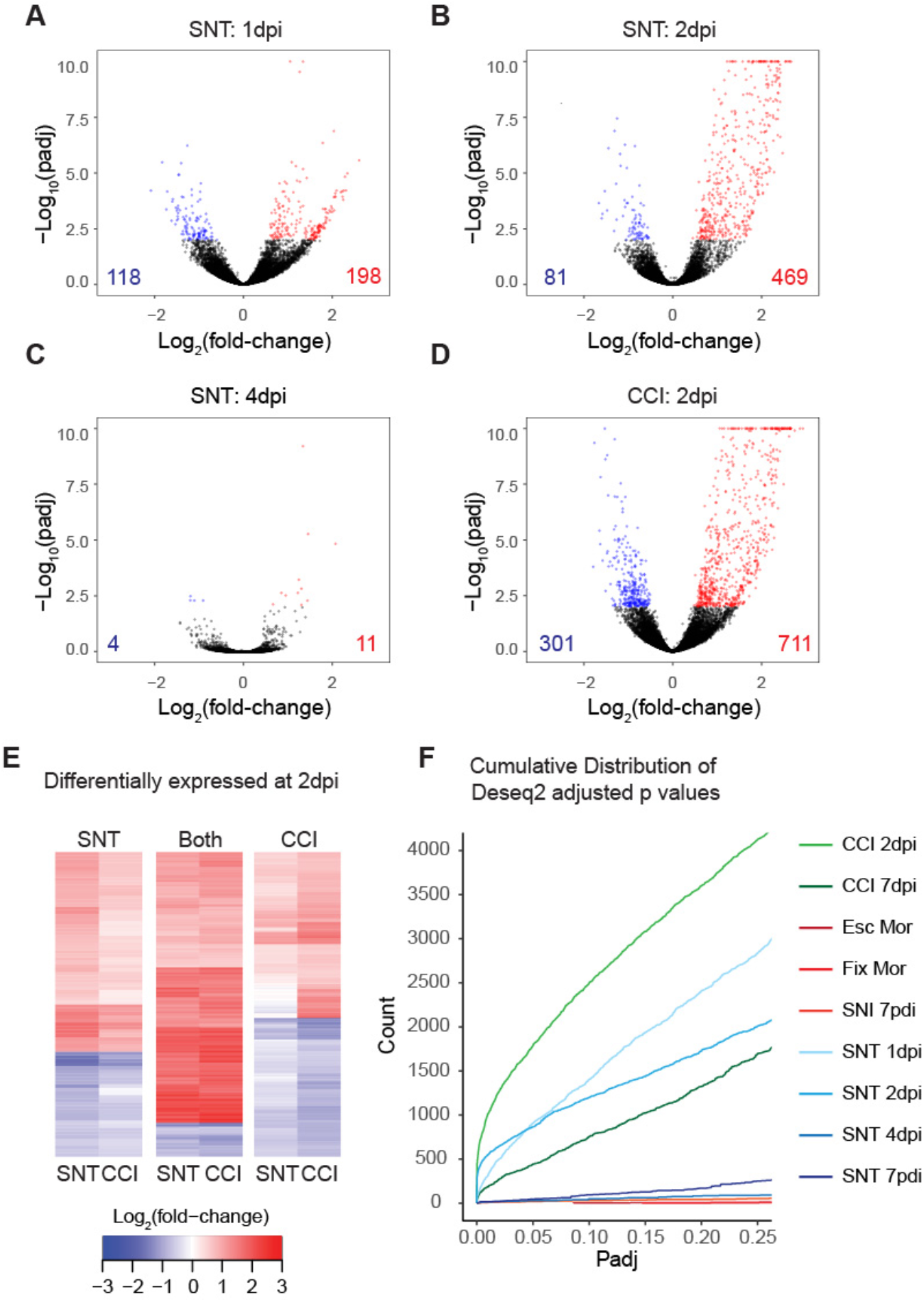
The greatest transcriptional variability occurs early after injury. (A-D) Volcano plots summarizing gene expression changes in microglia 1 (A), 2 (B), or 4(C) days post complete bilateral transection of 2(D) days after bilateral chronic constriction injury. Blue and red symbols indicate significantly down or upregulated genes from Deseq2, respectively (padj<0.01). Triangles indicate values outside the axis scale. The numbers in the lower corners indicate the total numbers of down (blue) and upregulated (red) genes at padj<0.01. (E) Heatmaps of genes differentially expressed 2 days post SNI only, both SNT and CCI 2 dpi, or 2 days post CCI only. Values are plotted as Log_2_(fold-change). (F) Cumulative distribution of padj values (≤0.25) from Deseq2 for all injury and treatment conditions.

**Figure 5:**
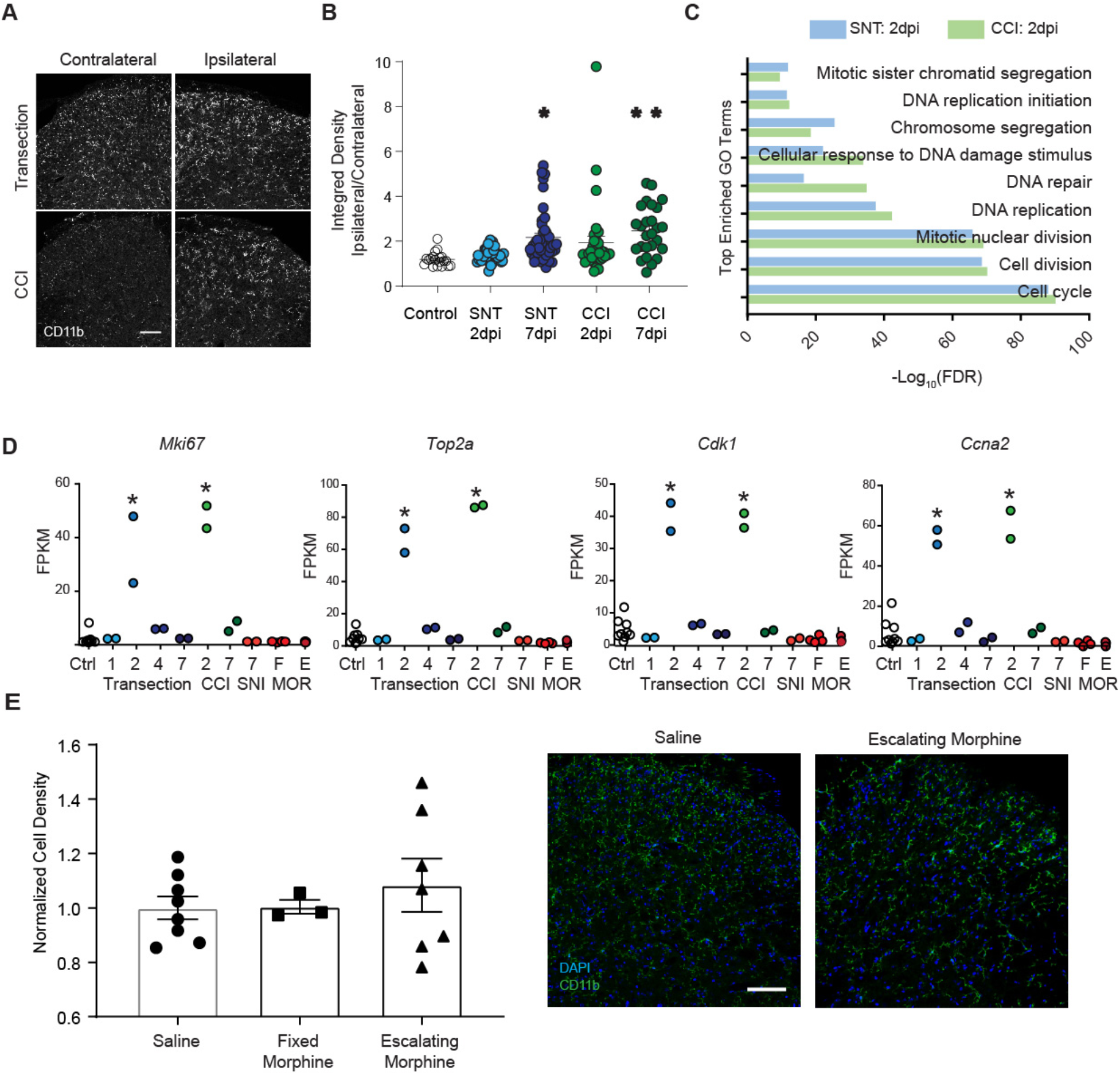
Proliferation drives microglial activation following PNI but not morphine treatment. (A) Example images of CD11b immunostaining in lumbar spinal cords 2 days after unilateral transection of CCI. (B) Quantification of spinal cord dorsal horn anti-CD11b immunoreactivity 2 and 7 days after transection or CCI compared to control spinal cord, as measured by a ratio of fluorescence integrated density in ipsilateral to contralateral dorsal horn. Ipsi/contra for control tissue was set randomly. (*p=0.104, **p=0.001) (C) Top enriched GO Terms associated with SNT (blue) or CCI (green) two days post injury. (D) Time course of normalized (FPKM) expression across time and injury for genes associated with proliferation. *indicates significantly differentially expressed with Deseq2 p<0.01 compared to relevant control groups. (E) Quantification of spinal cord microglia from saline or morphine treated mice. Data are plotted as normalized densities. Right: Representative images of saline and escalating morphine treated spinal cord dorsal horns showing CD11b/DAPI overlap. Scale bars = 100 μm.

Importantly, in comparison to SNT, SNI, or CCI chronic pain conditions, chronic opioid treatment did not produce a similar transcriptional signature (Figure 5I). The immune response genes detected following CCI showed no expression changes in either morphine dosing paradigm. These results confirm that opioid treatment induces a different microglial transcriptional response compared with nerve injury (particularly CCI), despite similar histological and hyperalgesic phenotypes. Strikingly, transcripts for BDNF, which is thought to be a key mediator for microglia contribution to both neuropathic pain and OIH ^11,15^, were undetectable in our transcriptional datasets.

### Proliferation drives transcriptional changes early after peripheral nerve injury

The unexpected disconnect between the dramatic histological changes and the limited transcriptional changes after peripheral nerve injury prompted us to conduct a time course study of microglial responses. We conducted this time course using the SNT model (rather than SNI) to maximize sensitivity for differentially expressed genes. We performed bilateral SNT and RNA-seq as described above at 7 dpi, but also collected tissue at 1, 2, and 4 days after injury. Differential expression analysis revealed extensive transcriptomic changes early after injury that were not detected at later time points (Figure 4 A,B). We identified 316 genes as differentially expressed 24 hours after injury (Figure 4A), and detected the greatest number of differentially expressed genes, 550, at 2 dpi (Figure 4B). In contrast, by 4 dpi, only 15 genes were differentially expressed (Figure 4C). We selected the time point with the greatest differential expression, 2dpi, for comparison to the CCI model. Two days after CCI, we detected 711 upregulated genes and 301 downregulated genes (Figure 4D), which represented the largest response from all groups. Additionally, we found extensive overlap between the lists of differentially expressed genes at 2dpi with SNT and CCI models, suggesting an analogous early microglial response between conditions (Figure 4E).

Overall, we found that the major transcriptional changes at 2 dpi had largely resolved at 4 and 7 days after SNT, and evolved into an inflammatory response by 7 days after CCI. We further confirmed that the increased number of differentially expressed gene calls at early time points after SNT or CCI was not a result of selecting a particular p-value cutoff, as the difference in the extent of changes were observed regardless of the significance cutoff selected (Figure 4F).

Histologically, microglia begin to take on an altered morphology and show increased CD11b immunoreactivity at this 2 dpi time point, but not to the extent that is observed at 7 dpi (Figure 5A and B), revealing a discrepancy between observed morphological changes and the extent of transcriptional changes during pain-associated microglial activation. To determine what classes of genes were upregulated at early time points in each condition we again used GO term analysis. We found that cell cycle and cell division markers were the clear driving force of differential expression between control and injury conditions at 2 dpi (Figure 5A), and the results were largely indistinguishable between SNT and CCI. This cellular response is clearly illustrated by the expression of canonical cell proliferation markers *Mki67, Top2a, Ccn2*, and *Cdk1*, which are almost exclusively expressed at 2 dpi in both conditions (Figure 5B). Collectively, these results demonstrate that similarities exist in the temporal structure of microglial responses across nerve injury models, with an early transformation dominated by proliferation markers that resolves within days. After resolution of the initial proliferative response, however, persistent alterations in microglial transcriptome are more limited, with only a few persistent changes universal to all injuries. These trends did not apply to chronic morphine treatment.

Our differential transcriptomic analysis thus suggests that while both opioids and PNI cause histological evidence of microglia activation, microglia only proliferate following PNI, arguing against common mechanisms for PNI- and opioid-induced hypersensitivity. To test this concept further, we treated additional cohorts of mice with saline or escalating morphine and measured microglia density by counting dorsal horn CD11b positive cells. We found no change in the number of microglia detected (Figure 5E), suggesting there is no microglial proliferation following chronic morphine treatment, confirming a fundamental divergence from PNI-induced microglial responses and hypersensitivity.

### Diversity of microglial transcriptional responses to insult and disease

Given that our analysis revealed a diversity of microglial transcriptional responses associated with hyperalgesic conditions, we next sought to compare these pain-associated responses to the microglial transcriptional signatures characteristic of other disease states. A meta-analysis across 336 purified myeloid transcription datasets identified modules of co-regulated genes characteristic of diverse disease or injury states ^31^. We utilized these modules to compare the transcriptomic responses of microglia to peripheral nerve injury or morphine treatment to those associated with proliferation, interferon-stimulation, LPS treatment, and neurodegenerative disease models (Figure 6). Importantly, and in agreement with our initial analysis, we found that genes implicated in microglial proliferation response are upregulated compared to control at 2 days post SNT or CCI, but are relatively unchanged at other time points and after morphine treatment (Figure 6A). Further supporting our interpretation that late time points are taking on an immune reactive phenotype, we found that genes associated with interferon stimulation were also associated with late time points following transection or CCI, such as the classic interferon-stimulated gene, *Ifit1* (Figure 6B). The stronger immune response induced by CCI was also reflected in the distribution of the LPS-treatment module, as illustrated by the complement component *C3* (Figure 6C). In stark contrast, morphine treatment did not induce an LPS-like signature of gene expression. While a minority of genes related to neurodegeneration were significantly upregulated in one or more of our manipulations, the proportion of genes from this module and the magnitude of these transcriptional changes were very limited compared to the changes associated with neurodegenerative models (Figure 6D). These observations strongly support the idea that microglial activation as detected by histological examination cannot be assumed to correspond to any stereotyped transcriptional state, but rather may reflect widely varying conditional responses.

**Figure 6:**
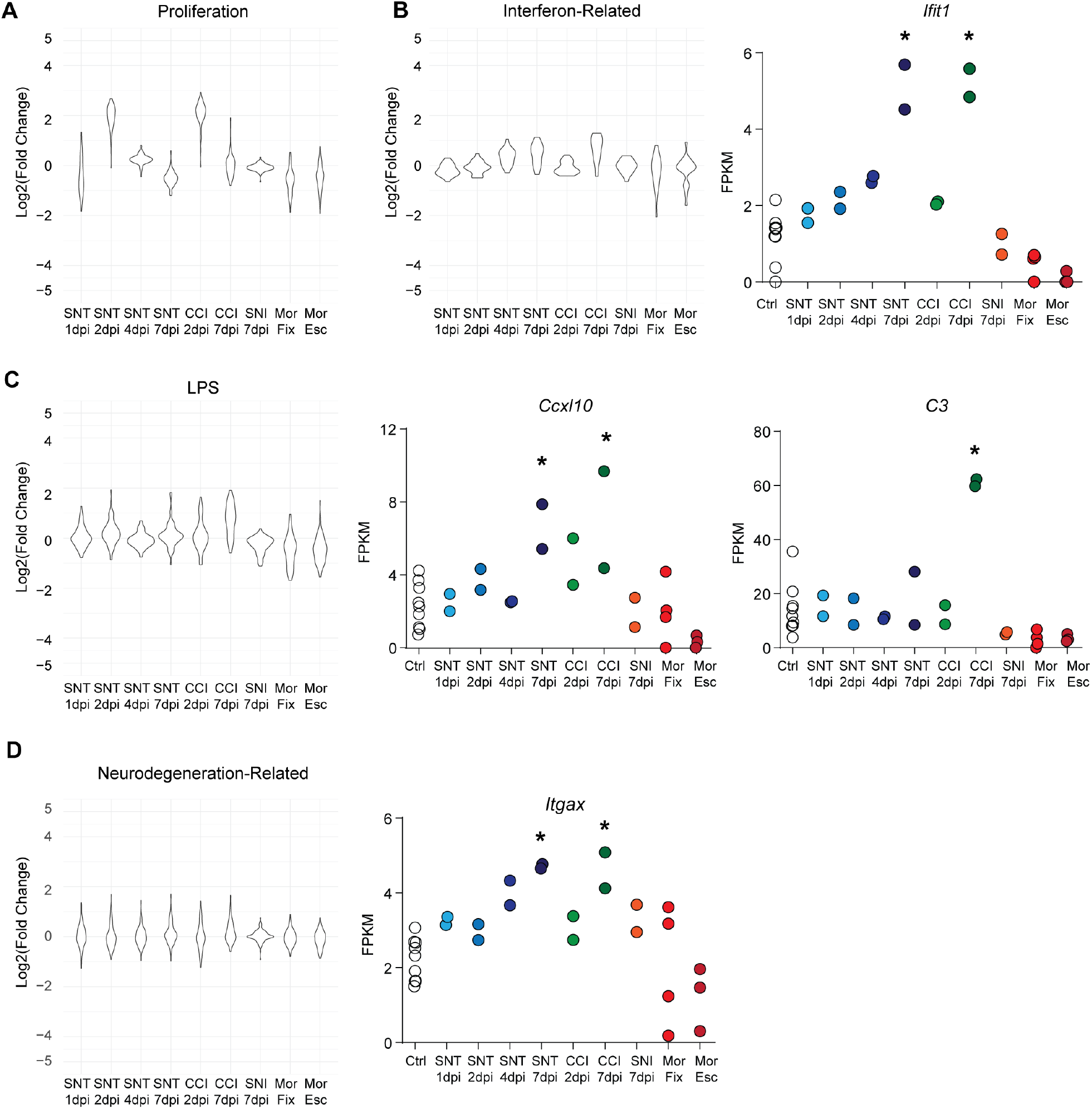
Diversity of microglial transcriptional responses Differential expression, displayed as Log2(fold-change) from DESeq2 output compared to control, for each gene in four modules selected from Friedman et al. 2018 across conditions tested with FPKM values plotted for example genes. Stars indicate significance from DESeq2 output.

Collectively, the transcriptional data presented here and the comparison with published datasets establishes the necessity of multimodal descriptions of activation states to resolve microglia molecular mechanisms of function, including following PNI- and opioid-induced hyperalgesia.

## Discussion

In this study, we provide a full transcriptional characterization of acutely isolated spinal cord microglia across multiple hyperalgesic states (chronic opioids, SNI, SNT, CCI,) with enhanced resolution of the progression of microglial reactivity. Microglial depletion after chronic morphine treatment reversed established hypersensitivity, supporting the importance of microglial function at this time point ^11^. Unexpectedly, we found no significant gene expression changes after opioid treatment. Previous studies suggested that microglia activation following morphine treatment occurs due to binding of morphine to MORs expressed by microglia ^11^. However, we were unable to detect MOR expression in spinal microglia using immunohistochemistry, *in situ* hybridization, or RNA-seq, and found that opioid-induced microglia activation is intact in MOR null mice ^10^. Further, we found no evidence of two other opioid receptors, kappa (*Oprk1*) and delta (*Oprd1*), ruling out morphine binding microglial DORs or KORs as a potential mechanism of microglial activation. An alternative mechanism of microglial activation implicates opioid engagement of microglial toll-like receptor 4 (TLR4) ^21,35,39^. TLR4 activation by LPS produces significant transcriptomic changes in vivo ^40,41^ and in culture (Figure 2), and we expected to detect similar transcriptional changes if morphine was indeed acting as a TLR4 agonist. Note that transcript for the TLR-4 co-signaling molecule MD-2, implicated the interaction of morphine and TLR4 ^35^, was not detected in our cultures. Nevertheless, the lack of transcriptional evidence of LPS-like signaling in response to morphine *in vivo* and *in vitro* argues against direct TLR4 activation, consistent with reports showing intact microglial activation or morphine tolerance in TLR4 null mice ^42,43^. Together, these data suggest that our transcriptional datasets include transcripts for other receptors than opioid receptor and TLRs that may act as sensors for microglia to detect either opioids, or secondary mediators released following opioid treatments.

To consider the possibility of biologically meaningful gene expression changes that were below our statistical threshold of detection, we further evaluated the trends in this dataset as they compared to the transcriptional response of spinal cord microglia to peripheral nerve injury. We show that following PNI microglia undergo a common transient, proliferation-driven transcriptional response. This signature is consistent with previous studies that used histological and imaging methods to identify microglia proliferation following PNI ^44–47^. We found greatest transcriptional variability 24-48 hours after injury, preceding maximum changes in histological markers of activation. Interestingly, depletion of microglia or their inhibition with minocycline is only effective at impacting pain hypersensitivity when introduced at these early proliferative time points ^48,49^. The minimal transcriptional changes detected at later time points may still be of substantial importance, as we also detected divergence between injury types 7dpi, with a strong inflammatory immune response only following CCI. Interestingly, CCI also selectively triggers an epineural inflammation compared to SNI, suggesting a commonality between peripheral and microglial immune responses to pronociceptive insults ^50^.

By performing a meta-analysis incorporating our datasets and those of Friedman et al 2017, we could further reveal the diversity of microglial responses. For example, even at time points with greatest transcriptional variability, PNI- and OIH-associated microglial transcriptional signatures differed from those associated with neurodegenerative conditions. Conversely, we do see similarity to LPS-induced transcriptional changes with the CCI model of PNI, but not with morphine treatment, as would be predicted by opioid activation of TLR4. These and other sequencing studies have aided the search for selective microglia markers and their activation state, though our findings suggest there may be no universal marker or signature across conditions.

There are of course mechanisms by which microglia can contribute to hyperalgesic states that do not rely on transcriptional changes. Epigenetic changes in microglia were identified following PNI, contributing to the long-term maintenance of neuropathic pain ^51. 52,53,54^. C1q and complement signaling are associated with synaptic pruning during development ^55^, and microglia may use similar complement-mediated mechanisms to remodel synapses in the dorsal horn after PNI or opioid treatment. Notably, despite being implicated in both neuropathic pain and OIH, we were unable to detect BDNF transcripts in the majority of our acutely isolated or cultured microglia samples, consistent with recently published microglial sequencing datasets ^51,56^. This adds complexity to interpreting studies in which genetic deletion of BDNF from microglia impacted not only pain but also learning and memory ^11,15,57^ and warrants further study to examine potential disconnect between transcript and protein presence. Alternatively, BDNF transcripts in astrocytes ^24^, dorsal root ganglion, ^58^ or spinal cord dorsal horn neurons ^59^ suggest that BDNF from non-microglial origins could alter dorsal horn excitability during neuropathic pain or OIH. Future proteomic studies may be required to fully resolve the mechanisms of BDNF production and action in the dorsal horn.

All experiments for this study were conducted in males. This decision was based on known differences in microglial responses between sexes, and the suggestion that peripheral immune cells, rather than central microglia, mediate hyperalgesia in females ^60,61^. This is not to suggest that microglia do not play a role in hyperalgesic conditions in females, simply that an entire adequate characterization based on sex is required on its own. Extensive differences in pain sensitivities have been documented between males and females, as well as differences in responsiveness to opioids ^62^.

Collectively, our data demonstrate how dynamic transcriptional responses of microglia can vary across disease models, including hyperalgesic states. Despite similarities in histological marker expression following peripheral nerve injuries or opioid exposure, microglia engage insult-specific mechanisms. We conclude that to battle the current Opioid Epidemic ^63^, both by limiting the side effects of opioids and discovering alternative treatments for pain, therapeutics targeting microglia will require the selective targeting of the pronociceptive mechanisms engaged in response to specific injuries, drugs or diseases rather than general inhibition of canonical inflammatory mediators.

## Acknowledgements

This work was supported by National Institutes of Health grants DA031777 and DA044481 (G.S.), the New York Stem Cell Foundation (G.S.), a Department of Defense Neurosensory Award MR130053 (G.S.), a Rita Allen Foundation and American Pain Society Award in Pain (G.S.), a DoD National Defense Science and Engineering Graduate (NDSEG) Fellowship (E.I.S.), NIH grant R37DA15043 (B.A.B.), and a Damon Runyon Cancer Research Foundation postdoctoral fellowship (C.J.B.). G.S. is a New York Stem Cell Foundation – Robertson Investigator. We thank Amaury François for his contributions to the development of this manuscript.

## Materials and Methods

### Animals

All procedures were approved by the Stanford University Administrative Panel on Laboratory Animal Care in accordance with the International Association for the Study of Pain. Male 8 - 12 week old C57Bl6/J mice from The Jackson Laboratory were housed 2-5 per cage and on a 12-hour light/dark cycle with ad lib access to food and water. Male Sprague-Dawley rats used for culture experiments only were obtained from Charles River, Hollister, CA.

### Drugs

Morphine sulfate (West-Ward NDC 0641-6070-01) was administered subcutaneously (s.c.) at 10 - 40 mg/kg (Figure 1A). Dilutions and control injections were prepared with 0.9% sodium chloride (Hospira NDC 0409-4888-10).

### Histology

#### Tissue Collection and Processing

Mice were anesthetized with sodium pentobarbital were transcardially perfused with phosphate-buffered saline (PBS) followed by 10% formaldehyde in PBS. Spinal cord was dissected and post-fixed in 10% formaldehyde for 4 hours, then cryoprotected overnight in 30% sucrose in PBS. Tissues were frozen in Tissue-Tek and sectioned using a cryostat (Leica). Tissue was sectioned at 40 μm and stored in PBS at 4°C and transferred to glycerol-based cryoprotectant solution for long term storage at −20°C.

#### Immunohistochemistry

Tissues were blocked for 1 hour in 0.1 M PBS with 0.3% Triton X-100 and 5% normal donkey serum (NDS). Primary and secondary antibodies were diluted in 0.1 M PBS with 0.3% Triton X-100 and 1% NDS. Sections were then incubated overnight at room temperature (RT) in primary antibody solution, washed in 0.1 M PBS with 0.3% Triton X-100 and 1% NDS 3×10min, incubated for 2 hours in secondary antibody (RT), and washed in 0.1 M PB for 3×10min. Sections were then mounted using Fluoromount-G on Fisherbrand Superfrost Plus microscope slides. Images were acquired with a Leica TCS SPE confocal microscope.

Primary antibodies: anti-CD11b, Abd Serotec # MCA711G (rat, 1:1,000). Alexa Fluor®-conjugated secondary antibodies were acquired from Invitrogen and Jackson Immunoresearch Labs.

#### Quantification of immunofluorescence and microglia number

For quantification of CD11b immunofluorescence in Figure 5A lumbar spinal cord tissue from all conditions analyzed was immunostained and imaged together. Single confocal plane images were taken of both dorsal horns. Using ImageJ, a region of interest was drawn around the dorsal horn to exclude image space without tissue. Ipsilateral and contralateral dorsal horns from each slice were processed together with the same threshold. CD11b integrated density was measured, and ipsilateral value was divided by contralateral for one measurement per slice. n=5-17 slices per animal, 2-3 animals per condition.

For counts of microglial density following chronic opioid treatment, mice were treated with saline or morphine (fixed or escalation dosing) and processed for CD11b immunohistochemistry as described above. Single confocal plane images of the dorsal horn were taken with DAPI labeling. Only clearly labeled microglia positive for both CD11b and DAPI were counted and normalized for tissue area. Because tissue was prepared and counted in two separate replicates by independent experimenters, results were normalized to the appropriate control tissue. Each treatment group and replicate had a minimum of 3 animals, with a minimum of 9 slices per animal.

#### Injury Models

Three peripheral nerve injuries were performed: complete transection of the sciatic nerve (SNT), chronic constriction injury of the sciatic nerve (CCI) and spared nerve injury (SNI) ^37,63^. Adult mice were anesthetized with isoflurane, and the right sciatic nerve was exposed at mid-thigh level. For CCI, two 5-0 silk sutures were loosely tied around the nerve about 2 mm apart. For SNT, a 2-mm portion of the nerve was removed. For SNI, only the tibial and common peroneal branches were cut. The muscle was pinched closed and the skin was sealed with tissue adhesive (Vetbond). For RNA sequencing the procedure was repeated on the left side and spinal cords were collected 1, 2, 4, and 7 days after transection, 2 and 7 days after CCI, and 7 days after SNI. For histology, the injury was only performed on the right side.

#### Microglia RNA library construction and sequencing

Mice were perfused with ice cold PBS and caudal lumbar spinal cords were isolated. Caudal lumbar segments from 4 to 15 mice per group were pooled to obtain enough material to ensure tight correlation between replicates. Tissue was mechanically dissociated in HBSS (Gibco) containing 0.5% Glucose, 15 mM HEPES pH 7.5, and 125 U/mL DNaseI then processed with MACS myelin removal (Myelin Removal Beads II, Miltenyi) followed by CD11b selection (CD11b Microbeads, Miltenyi) according to the manufacturer’s instructions, except all centrifugation steps were shortened to 30 s at 10,000 rpm.

Total RNA was extracted from CD11b-positive cells using the RNeasy Micro Kit (Qiagen). Libraries were prepared using Smart-Seq2 ^64^ and modified for sequencing using the Nextera XT DNA Sample Preparation Kit (Illumina) with 300 pg of cDNA as input material. Libraries were sequenced using the Illumina Nextseq or Miseq to obtain 75 bp paired-end reads. At least two libraries for each condition were prepared and sequenced independently for an average of 10.1 million reads per group (range of 1.8-28.6M).

Reads were mapped to the UCSC mouse reference genome mm10 via the Galaxy platform (http://usegalaxy.org) using HISAT2 ^65^ version 2.0.5.1. FPKM (fragments per kilobase of transcript sequence per million mapped fragments) values were obtained using Cufflinks ^66^ version 2.2.1. Reads were quantified using featureCounts version 1.4.6.p5 ^67^. Differential analysis was conducted in R with DeSeq2 ^68^. A stringent significance cutoff was set using the adjusted p value <0.01 to account for low replicates in some conditions. No statistical analysis was performed on FPKM values, stars in figures illustrating FPKM values are derived from Deseq2 results. GO term analysis was conducted by creating individual lists of up and down regulated genes in each condition with padj<0.01 from Deseq2 output and evaluated using DAVID ^69,70^. Initial analysis of *Oprm1* expression from a subset of CCI, SNT, and morphine datasets was reported in our previous work ^10^. All sequencing data are available at GEO GSE117321.

#### Microglia culture and stimulation

Rat microglia were isolated and cultured as described ^36^. Briefly, brains were extracted from perfused male 21 day old rats and the cerebellum of each brain was discarded. Brains were mechanically dissociated to a single-cell suspension and passed through a density gradient to remove myelin and cellular debris. Myeloid cells were isolated by CD11b-immunopanning and plated in serum-free medium (DMEM/F12 containing 100 units/mL penicillin, 100 ug/mL streptomycin, 2mM glutamine, 5 ug/ml N-acetyl cysteine, 5 ug/ml insulin, 100 ug/mL apo-transferrin, 100 ng/mL sodium selenite, 0.1 ug/mL oleic acid, 0.001 ug/mL gondoic acid, 1 ug/mL heparan sulfate, 1.5 ug/mL ovine wool cholesterol, 100 ng/mL murine IL-34, and 2 ng/mL human TGF-β2). Cells were maintained for 5 days with 50% medium change on day 3. After 5 days, cell were stimulated with TLR ligands LPS (1 ng/mL), PAM3CSK4 (1 ng/mL), or zymosan (100 ug/mL) or morphine (1 uM or 10 uM) by introducing the drug in a 50% medium change.

#### Cultured cell gene expression analysis

For gene expression experiments, cells were incubated with morphine for 4 or 24 hours or TLR agonist for 24 hours, then washed twice before lysis and RNA purification using the RNeasy Micro kit (Qiagen). For RNA-seq experiments, samples were prepared and analyzed as described above for freshly isolated mouse samples except data was mapped using the rat genome annotation rn6. For QPCR experiments, RNA was reverse-transcribed using a High-Capacity RNA-to-cDNA Kit (ThermoFisher Scientific) and used as template for SYBR Green PCR Master Mix (ThermoFisher Scientific) reactions (20 uL volume) with previously published primers ^36^. CT values were registered using noiseband thresholding, and melting curves were checked to ensure formation of a single product within the expected size range. ΔCT values were calculated relative to the housekeeping gene RPLP0. Consistent ΔCT values of a second housekeeping gene, EEF1A1, were measured across samples. All CT values were detected before completion of the 38th cycle.

